# Plasminogen Activator Inhibitor-1 is internalized by endothelial cells via macropinocytosis to evade degradation

**DOI:** 10.1101/2024.09.26.615054

**Authors:** Molly McAdow, Gjina Ahmetaj, Rachel Hauschel, Anne Eichmann, William Sessa

## Abstract

Plasminogen activator inhibitor1 (PAI1) promotes hemostasis and is a biomarker of cardiovascular disease but accumulating evidence supports a role for PAI1 in intracellular biology. Recently, we found that PAI1 directly inhibits endothelial nitric oxide synthase (eNOS) and that exogenous PAI1 is internalized and traffics to eNOS. However, prior work has demonstrated that when PAI1 is internalized via LRP1 and uPAR on the cell surface and is internalized and degraded. Our objective was to identify the mechanism by which PAI1 is internalized and evades degradation. Here, we show that PAI1 is internalized by endothelial cells in an energy-dependent manner and persists in the cell for at least 6 hours. Entry is independent of LRP1, uPAR, or clathrin-mediated endocytosis. It is internalized in large vesicles, is inhibited by amiloride and nocodazole, and is partially dependent on CDC42, consistent with macropinocytosis. We propose that following internalization by macropinocytosis, PAI1 evades degradation, permitting eNOS inhibition.

## Introduction

Plasminogen activator inhibitor 1 (PAI1) is a secreted serine protease inhibitor that is elevated in numerous disease states including preeclampsia [1], type 2 diabetes [2], and cardiovascular disease [3, 4]. PAI1 inhibits tissue- and urokinase-type plasminogen activators (tPA and uPA, respectively), thereby stabilizing fibrin clots [5] and promoting vascular fibrosis [6]. A canonical signal peptide at the N-terminus of PAI1 drives its secretion to the extracellular space, which is up-regulated by TNFα, TGFβ, Angiotensin II, and hypoxia [7–10].

PAI1 is a suicide inhibitor of its target proteases. It is secreted in an active conformation, which has a reactive center loop with an exposed “bait sequence” for tPA/uPA. When tPA/uPA cleaves the bait sequence, it triggers a conformational change in PAI1 in which the reactive center loop folds inward, trapping tPA/uPA [5]. The active conformation is thermodynamically unstable. *In vitro*, the active conformation has a short half-life and converts to a latent conformation in which the reactive center loop is not accessible. Prior studies on the fate of extracellular PAI1 have focused on internalization of PAI1 when in complex with tPA/uPA. By monitoring ^125^I-labelled uPA complexed with PAI-1, it was found that the complex binds urokinase-type plasminogen activator receptor (uPAR) and lipoprotein related peptide 1 (LRP1), which triggers internalization and degradation of the radio-labeled substrate while LRP1 and uPAR are recycled to the cell surface [11–13].

In addition to its canonical role in the extracellular space, accumulating evidence suggests that PAI1 also alters intracellular signaling. PAI1 inhibits PI3K/Akt growth signaling, at least partially dependent on its interaction with LRP1 on the cell surface [14, 15], and interaction between PAI1 and LRP1 on the cell surface may provoke Jak/STAT activation, promoting endothelial migration [16]. Direct intracellular activity by PAI1 has been demonstrated by several groups. Within the Golgi, PAI1 inhibits furin proprotein convertase, leading to reduction in maturation of insulin receptor and ADAM17 [17]. Our group recently showed that exogenous PAI-1 directly interacts with and inhibits endothelial nitric oxide synthase (eNOS) [18]. Moreover, using a C-terminal hemagglutinin tag, it was found that exogenous PAI1 is internalized and traffics to eNOS. Most recently, PAI1 was observed within the nucleus and found to bind DNA via chromatin immunoprecipitation, suggesting that PAI1 may be a transcription factor [19]. Together, these finding demonstrate intracellular effects of PAI1, a secreted protein, and suggest that PAI1 can escape degradation upon endocytosis. However, the mechanism by which PAI1 is internalized yet evades degradation is not known.

Our objective was to determine the mechanism through which PAI1 is internalized by endothelial cells and escapes degradation. Here, we demonstrate that exogenous PAI1 is taken up by cultured endothelial cells through macropinocytosis that does not require interaction with uPAR or LRP1 and can escape degradation.

## Methods

### Cell culture

Ea.hy926 cell line was purchased form the American Type Culture Collection (CRL-2922) and grown in Dulbecco’s Modified Eagle Medium (DMEM) (Gibco) containing 10% fetal bovine serum (FBS), penicillin (100 U/mL), streptomycin (0.1 mg/mL), glutamine (2mM), and hypoxanthine-aminopterin-thymidine (HAT) as previously described [18]. Primary human umbilical vein endothelial cells (HUVECs) were obtained from the Yale University Vascular Biology and Therapeutics Core facility, plated on 0.1% gelatin-coated dishes in Endothelial Cell Growth Medium (Lonza) and used at passage 4. COS7 cells were cultured in DMEM supplemented with 10% FBS, penicillin (100 U/mL), streptomycin (0.1mg/mL), and glutamine (2mM).

### Transfection

For experiments with silencing RNA (siRNA), endothelial cells were grown to 70% confluence and transfected with 20nM siRNA targeting LRP1 (Dharmacon), uPAR (Dharmacon), clathrin heavy chain (ThermoFisher), caveolin 1 (Dharmacon), dynamin 2 (Qiagen), CDC42 (Dharmacon), Rac1 (Dharmacon), or control non-targeting pool siRNA (Dharmacon) using Lipofectamine RNAiMax (ThermoFisher Scientific) in Opti-MEM medium (ThermoFisher Scientific). After 8 hours, the media was replaced with complete media containing 10% FBS as above. Experiments were conducted after 48 hours of RNA silencing.

A construct for over-expression of PAI-1 with a C-terminal hemagglutinin (HA) tag was previously generated [18]. COS7 cells were transfected with 1 µg plasmid DNA (pcDNA3 empty vector or PAI1-HA) per 10mm culture dish using Lipofectamine 2000 (ThermoFisher Scientific) in Opti-MEM medium as previously described. After 8 hours, the media was replaced with complete media containing 10% FBS as described above. After 48 hours of incubation, PAI1-HA or pcDNA3 enriched spent media was collected and strained through a 40µm cell-strainer. Spent media was used fresh or frozen at -80°C for later use.

### PAI1-HA Endocytosis

Ea.hy926 or HUVECs were grown to 90% confluence in 6 well dishes. PAI1-HA enriched media was added to each well for 2 hours unless otherwise stated and cells were incubated at 37°C. In some experiments, cells were pre-treated with 5-(N-Ethyl-N-isopropyl) amiloride (Sigma) at the concentration listed or nocodazole (10µM, Sigma) for 30 minutes prior to addition of enriched media. Cells were then washed with ice-cold PBS. In some experiments, cells were also washed with acidic wash buffer (25mM glycine pH4), and cell lysates were collected in lysis buffer with the aid of cell scrapers as previously described [18].

### Immunoblotting

Protein samples were sonicated, and cell debris removed via centrifugation. Protein concentration was determined via the Bradford Assay. Samples were boiled in SDS loading buffer and separated by sodium dodecyl sulfate polyacrylamide gel electrophoresis (SDS-PAGE) and then transferred to 0.45 µm nitrocellulose membranes (Bio-Rad). The following primary antibodies were used: HSP90 (Santa Cruz), HA (Roche), LRP1 (Cell Signaling, 64099S), uPAR (Abcam), Clathrin heavy chain (BD), caveolin 1 (BD Biosciences), Dynamin 2 (Abcam), CDC42 (Abcam), Rac1 (EMD Millipore) at 1:1,000 dilution. The appropriate Li-COR secondary antibodies were used at 1:10,000. The Li-COR Odyssey Infrared Imaging System was used to detect and quantify immunoblots. The PAI1-HA signal was normalized to HSP90. For internalization time course, PAI1-HA uptake was band intensity normalized to HSP90. For all other western blots, HA signal at 2 hours was used as the reference and the percent PAI1-HA uptake was calculated as the percent difference between signal in the treatment group compared to the control, divided by the control.

### Immunofluorescence Microscopy

HUVECs grown on coverslips coated with 0.1% gelatin were treated with media enriched for PAI1-HA for 2 hours or as otherwise stated. In some experiments, rhodamine-conjugated dextran (Invitrogen) (1.5mg/mL) or Transferrin-tetramethylrhodamine (Invitrogen) (50µg/mL) were added to cells in pcDNA3-enriched media or PAI1-HA-enriched media. Cells were washed, fixed with ice-cold methanol at -20°C for 10 minutes, and incubated with blocking buffer (PBS containing 3% bovine serum albumin) for 1 hour at 37°C. Cells were then incubated with primary antibody diluted 1:100 in blocking buffer and incubated overnight at 4°C. Slides were washed and incubated with appropriate AlexaFluor-conjugated antibody (Life Technologies) secondary antibody diluted 1:1000 for 1 hour. Hoechst nuclear antibody stain was applied, and coverslips were mounted on slides with mounting media. Samples were visualized using LeicaSP5 confocal microscope. To determine PAI1-HA internalization, 20-40 random fields of view containing endothelial cells with normal morphology were captured with the same exposure time and gain settings. Each image was analyzed in ImageJ. For each field of view, the number of cells was counted by nuclei, and there were at least 200 cells counted per experiment. The color balance for the 488 channel (PAI1-HA) was increased to detect the cell boundary by autofluorescence, and the cell borders were demarcated. The color balance was then reset, and the maximum intensity within the cell was measured.

### Statistical Analysis

Statistical analysis was performed in GraphPad Prism 9. Results of Western blot analyses are presented as means ± SEM. Quantitative analysis of PAI1-HA internalization by immunofluorescence microscopy was found to follow a non-normal distribution; therefore, results are depicted as median ± interquartile range. All experiments in which more than one comparison were tested were analyzed by two-way analysis of variance (ANOVA) followed by Dunnett’s multiple comparisons test. Differences between two groups were compared by unpaired Student’s *t* test. *P* ≤ 0.05 was considered significant.

## Results

### Exogenous PAI-1 is endocytosed and persists in cultured endothelial cells

To confirm that exogenous PAI-1 can be endocytosed by endothelial cells yet evade degradation, PAI-1 with a C-terminal hemagglutinin (HA) tag (PAI1-HA) was added to human umbilical vein endothelial cells (HUVECs) in culture. Cells were washed and lysed, and quantitative Western blotting to the HA tag was performed to assess PAI1-HA internalization by endothelial cells. PAI1-HA is taken up and persists over at least 6 hours in HUVECs (Figure 1A) and Ea.hy926 cells (Figure 1B). PAI1-HA enriched media was added for 2 hours, switched to normal medium, and cell lysates were collected at time intervals, which demonstrated that PAI1-HA evades degradation over at least 6 hours (Figure 1C). To confirm that PAI1-HA is endocytosed and not simply associated with the cell surface, PAI1-HA enriched media was added to endothelial cells for 2 hours followed by washing with PBS or acidic wash buffer. PAI1-HA is not removed by acid wash (Figure 1D). When PAI1-HA was added to endothelial cells on ice for 2 hours, minimal PAI1-HA was internalized compared to incubation at 37°C (Figure 1E). Internalization was also confirmed by immunofluorescence microscopy (IF) with an antibody that recognizes the HA tag to visualize the fate of exogenous PAI1-HA (Figure 1F). Intensity of PAI1-HA signal within the cell increased over time. These findings suggest that PAI-1 is internalized by endothelial cells through an energy-dependent process and evades degradation.

**Figure 1:**
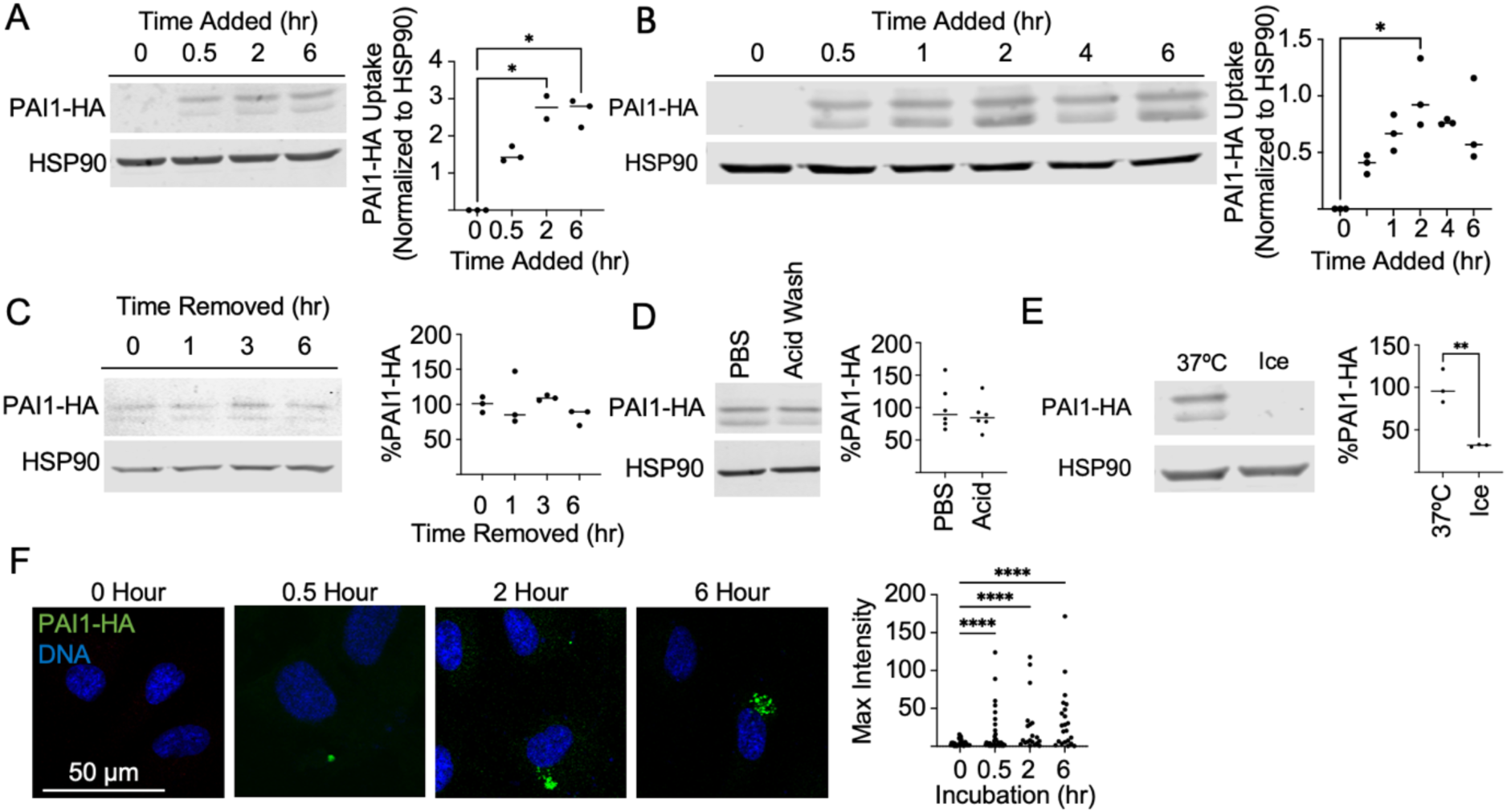
Exogenous PAI-1 is endocytosed by endothelial cells and persists over time. A) Spent media enriched with PAI-1 with a C-terminal hemagglutinin (HA) tag was added to primary human umbilical vein endothelial cells (HUVECs) for the specified durations at 37°C. Cells were washed, lysed, and lysates run on SDS-PAGE. Internalized PAI-1 was detected by immunoblotting toward the HA tag. Bar graph displays mean ± SEM of triplicate experiments, normalized to HSP90 loading control. B) Experiment conducted as in A but using Ea.hy926 cells. C) PAI1-HA spent media was added to HUVECs for 2 hours, then replaced with growth media for the specified durations and persistent PAI1-HA was assessed as described above. Bar graph displays mean ± SEM of triplicate experiments, normalized to HSP90 loading control. D) To confirm that PAI1-HA was actively internalized, rather than surface associated, after exposure to PAI1-HA for 2 hours, Ea.hy926 endothelial cells were washed with acidic wash buffer prior to lysis and run on SDS-PAGE. E) To confirm that this internalization was an active process, cells were incubated on ice for 2 hours with PAI1-HA enriched media and run on SDS-PAGE. F) Experiment conducted as described in A. Cells were then fixed and probed with anti-HA antibody followed by fluorescently conjugated secondary antibody and Hoechst nuclear stain and visualized by confocal microscopy. PAI1-HA internalization was quantified as maximum intensity of HA signal within the cell. All experiments were conducted at least three independent times for reproducibility. Statistical significance: * denotes *P* < 0.05, ** denotes *P* < 0.01, *** denotes *P* < 0.001.

### PAI1-HA uptake is independent of classical endocytic mechanisms

Previous work has demonstrated that uPA/PAI-1 internalization is dependent on LRP1 and uPAR [12, 20, 21]. To test whether exogenous PAI-1 that persists in endothelial cells is taken up by this mechanism, we used small interfering RNA (siRNA) to silence LRP1 or uPAR. Efficient silencing of LRP1 and uPAR was confirmed by Western blot 48 hours after transfection (Figure 2A). PAI1-HA was added for 2 hours, and internalized PAI1-HA was visualized and quantified by Western blotting. Silencing LRP1 or uPAR did not affect PAI1-HA endocytosis (Figure 2A). No defect in PAI1-HA internalization was detected when LRP1 or uPAR were silenced as visualized by IF (Figure 2B).

**Figure 2:**
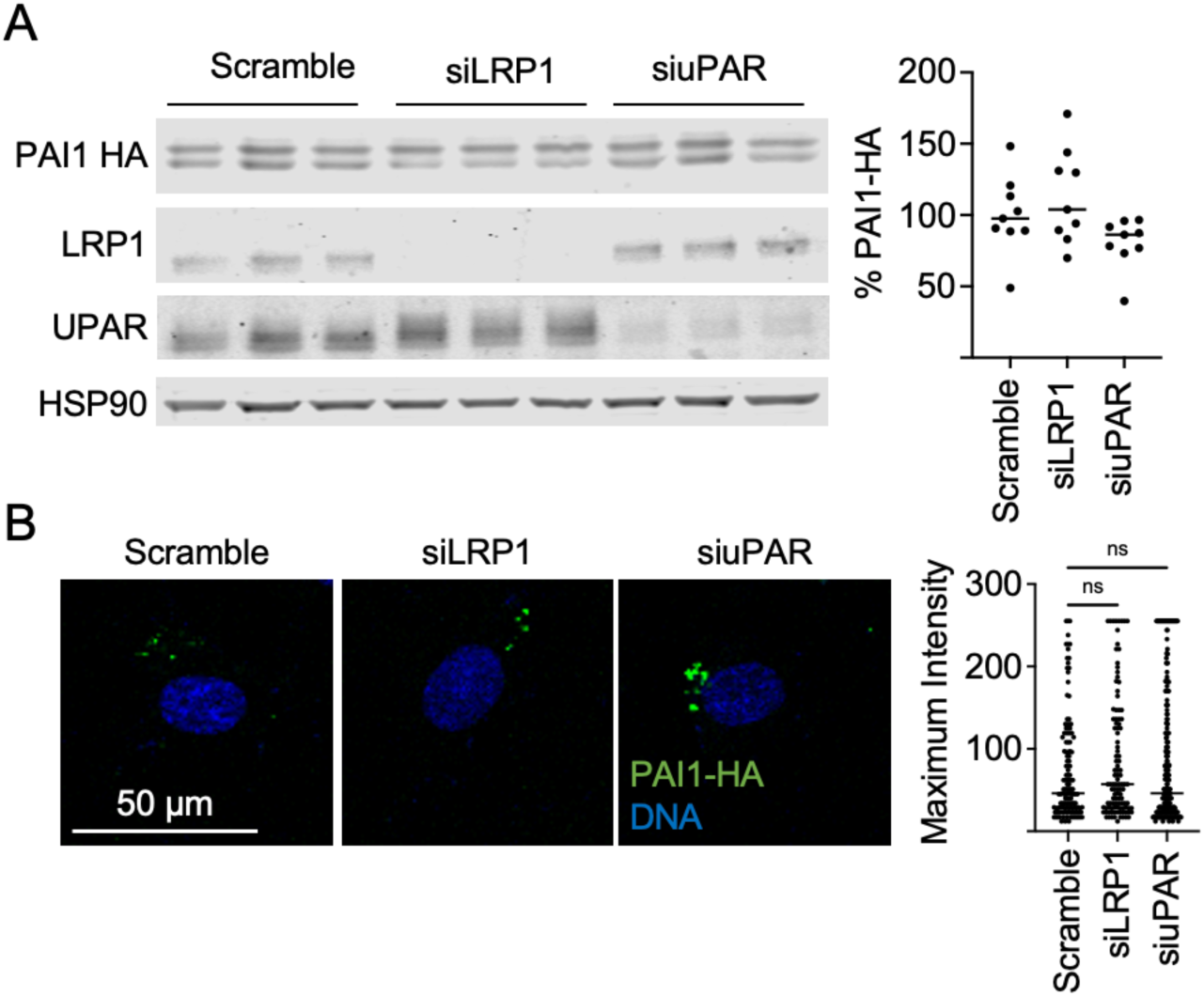
PAI1-HA persistence is not dependent on LRP1 or uPAR. A) siRNA was used to silence LRP1 and uPAR in Ea.hy926 cells. After 48 hours, cells were exposed to PAI1-HA spent media for 2 hours prior to lysis. Internalized PAI-1 was detected by immunoblotting toward the HA tag. Bar graph displays mean ± SEM of three biological replicate experiments from A, normalized to HSP90 loading control. B) Representative images of PAI1-HA puncta in cells treated with control siRNA or siRNA targeted LRP1 or uPAR. Graph depicts quantification of maximum intensity within each cell measured. Line depicts median.

Two well-described endocytic pathways employed by endothelial cells are clathrin-mediated endocytosis and caveolae, both of which use the GTPase Dynamin2 to pinch off endocytic vesicles from the plasma membrane [22]. To investigate whether clathrin-mediated endocytosis or caveolae are used for PAI1-HA endocytosis, siRNA was used to silence clathrin heavy chain, caveolin-1, or dynamin2 prior to PAI1-HA addition. Despite efficient silencing of clathrin heavy chain, caveolin-1, and dynamin2, there was no decrease in PAI1-HA internalization as measured by semi-quantitative Western blotting (Figure 3 A). To confirm that silencing of these endocytic mediators did not leave PAI1-HA localized to the cell surface, dynamin2 was silenced, cells were incubated with PAI1-HA for 2 hours, and cells were washed with acidic buffer prior to lysis. Acid wash did not reduce PAI1-HA uptake in these conditions (Figure 3B). Finally, silencing Dynamin2 did not affect PAI1-HA internalization as visualized by IF (Figure 3C). These results suggest that PAI1-HA internalization does not depend on either clathrin-mediated endocytosis or caveolae, two well described mechanisms of endocytosis in endothelial cells.

**Figure 3:**
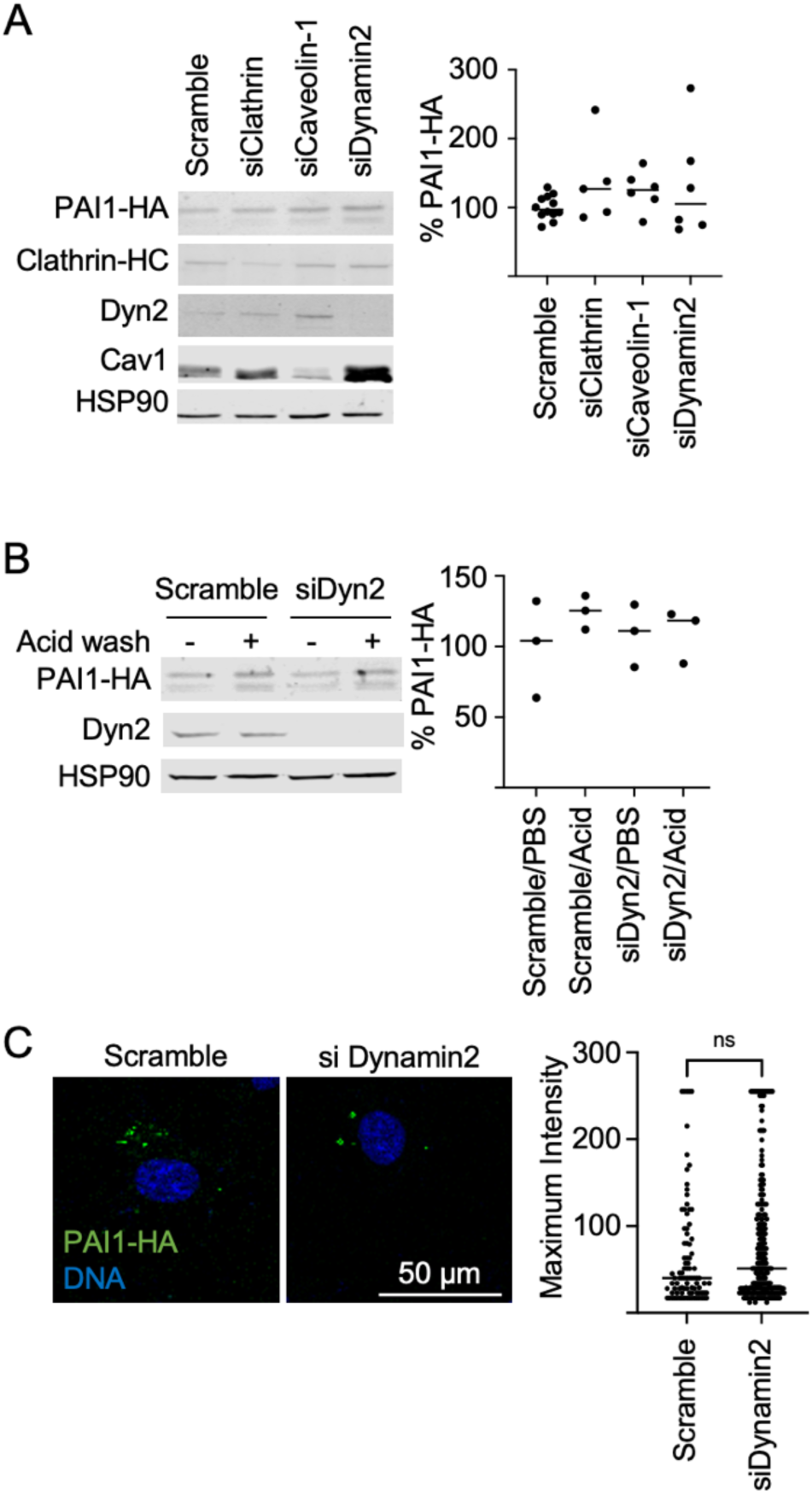
PAI1-HA endocytosis by endothelial cells is clathrin- and caveolin-independent. A) Ea.hy926 cells were transfected with siRNA targeting clathrin heavy chain, caveolin-1, or Dynamin2 for 48 hours. PAI1-HA spent media was added for 2 hours prior to lysis. Graph displays mean ± SEM of triplicate experiments, normalized to HSP90 loading control. B) Ea.hy926 cells were transfected with siRNA targeted dynamin2 for 48 hours and incubated with PAI1-HA enriched media for 2 hours. Cells were washed with PBS or acidic wash buffer prior to lysis. Graph displays mean ± SEM of triplicate experiments. C) Representative images of PAI1-HA puncta in cells treated with control siRNA or siRNA targeting Dynamin 2. Graph depicts quantification of maximum intensity within each cell measured. Line depicts median. One-way ANOVA or t-tests were used as appropriate.

### PAI1-HA uptake has features of macropinocytosis

Endothelial cells use macropinocytosis for signaling of fibroblast growth factor (FGF) and vascular endothelial growth factor (VEGF) [23–25]. Amiloride is a Na^+^/H^+^ exchange inhibitors that inhibit macropinocytosis [26]. To investigate whether amiloride inhibits PAI1-HA uptake, endothelial cells were pre-treated with amiloride for 30 minutes prior to addition of PAI1-HA enriched media. Amiloride blocked PAI1-HA uptake (Figure 4A). Pre-treatment of cells with nocodazole, an inhibitor of microtubule formation and macropinocytosis [27], led to approximately 50% less PAI1-HA internalization compared to vehicle-treated cells (Figure 4B). To confirm that PAI1-HA internalization was blocked in the presence of amiloride, cells treated with amiloride prior to adding PAI1-HA were visualized by IF microscopy. PAI1-HA internalization was reduced after amiloride treatment (Fig 4C). These findings suggest that PAI1-HA uptake may occur via macropinocytosis.

**Figure 4:**
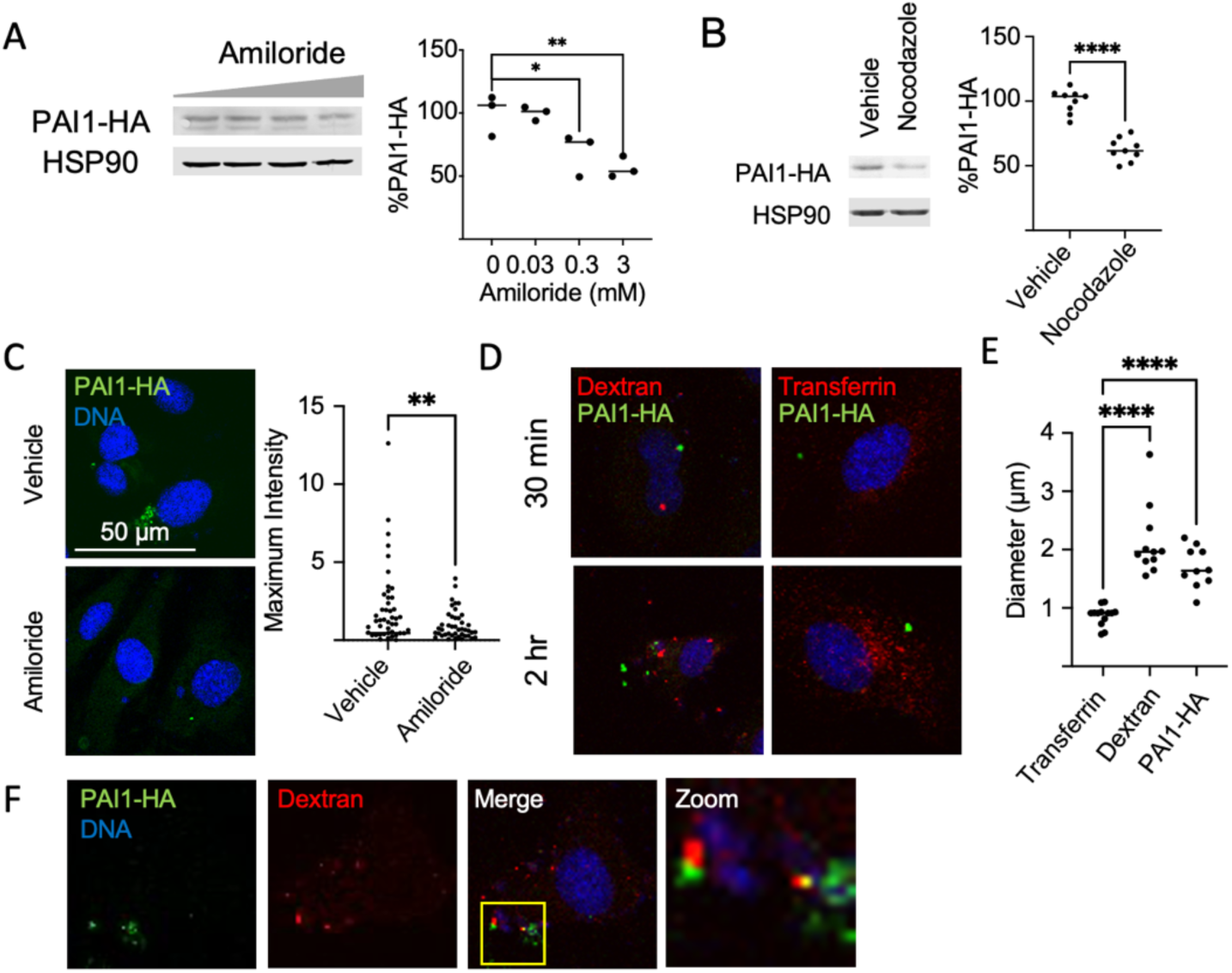
PAI1-HA endocytosis is amiloride-sensitive. A) HUVECs were pre-treated with increasing concentrations of amiloride for 30 minutes prior to addition of PAI1-HA spent media for 2 hours. Bar graph displays mean ± SEM of triplicate experiments from B, normalized to HSP90 loading control. B) Cells were pre-treated with nocodazole 10uM or vehicle for 30 minutes prior to addition of PAI1-HA spent media for two hours. Bar graph displays mean ± SEM of triplicate experiment normalized to HSP90 loading control. C) Cells on glass coverslips were treated with amiloride prior to PAI1-HA spent media for 2 hours. After treatment and fixation, cells were probed with anti-HA antibody followed by fluorescently conjugated secondary antibody and Hoechst nuclear stain. Graph depicts quantification of maximum intensity within each cell measured. D) PAI1-HA spent media was co-incubated with fluorescently labeled transferrin or 70kDa dextran for 30 minutes and 2 hours and visualized by IF microscopy. E) Fluorescent puncta within the cells were measured on the confocal microscope. Bar depicts median size. F) PAI1-HA spent media was co-incubated with fluorescently conjugated dextran for 2 hours and visualized by confocal microscopy. Representative images of two cells demonstrating colocalization of PAI1-HA and dextran. PAI1HA and dextran colocalized. Statistical significance: * denotes *P* < 0.05, ** denotes *P* < 0.01, **** denotes *P* < 0.0001. All experiments were conducted at least two independent times for reproducibility.

Macropinocytosis is a non-specific process in which membrane ruffles are made by actin polymerization and invagination of the extracellular fluid. Ea.hy926 cells were co-incubated with PAI1-HA and fluorescently conjugated transferrin, a cargo of clathrin-mediated endocytosis [22] or fluorescently conjugated 70kDa dextran, a classic marker of macropinocytosis [28]. Whereas transferrin is rapidly internalized within 30 minutes, PAI1HA and dextran are taken up more slowly (Figure 4D). Macropinosomes are relatively large endocytic vesicles compared with clathrin mediated endocytosis [22, 25, 29]. PAI1-HA was found in larger vesicles measuring 1.70 µm (standard deviation 0.35), similar to dextran-containing vesicles (mean 2.13 µm, standard deviation 0.60) compared to transferrin (mean 0.8673 µm, SD 0.17), *p*<0.0001 (Figure 4E). In some but not all vesicles, colocalization of PAI1-HA and dextran was observed (Figure 4F). There was no colocalization of PAI1-HA and transferrin. These findings are consistent with PAI1-HA internalization being via a macropinocytosis-like mechanism.

### GTPase CDC42 contributes to PAI1-HA internalization

Macropinocytosis is a process of non-specifically internalizing the extracellular fluid phase by polymerizing actin to form membrane ruffles, which are then closed off into pinocytic vesicles through activity of small GTPases including Rac1 and CDC42 and their effector kinase, p21 associated kinase (Pak1) [26, 29–33]. To investigate whether PAI1-HA uptake is dependent on Rac1 or CDC42, endothelial cells were transfected with specific siRNA for 48 hours prior to addition of PAI1-HA. Silencing CDC42 led to a small decrease in PAI1-HA internalization (Figure 5A). Silencing Rac1 led to no appreciable difference in PAI1-HA internalization at 2 hours as detected by Western blot and there was possibly increased abundance noted in some experiments (Figure 5B). After silencing macropinocytosis effector genes, cells were exposed to PAI1-HA for 2 hours and uptake visualized by immunofluorescence microscopy. Inconsistent with the Western blot results, there was no difference in PAI1-HA internalization after silencing CDC42 or Rac1 (Figure 5C), though we note that large accumulations of PAI1-HA were not observed when CDC42 was silenced. Consistent with sensitivity to amiloride and nocodazole, these findings support the hypothesis that PAI1-HA is internalized by macropinocytosis.

**Figure 5:**
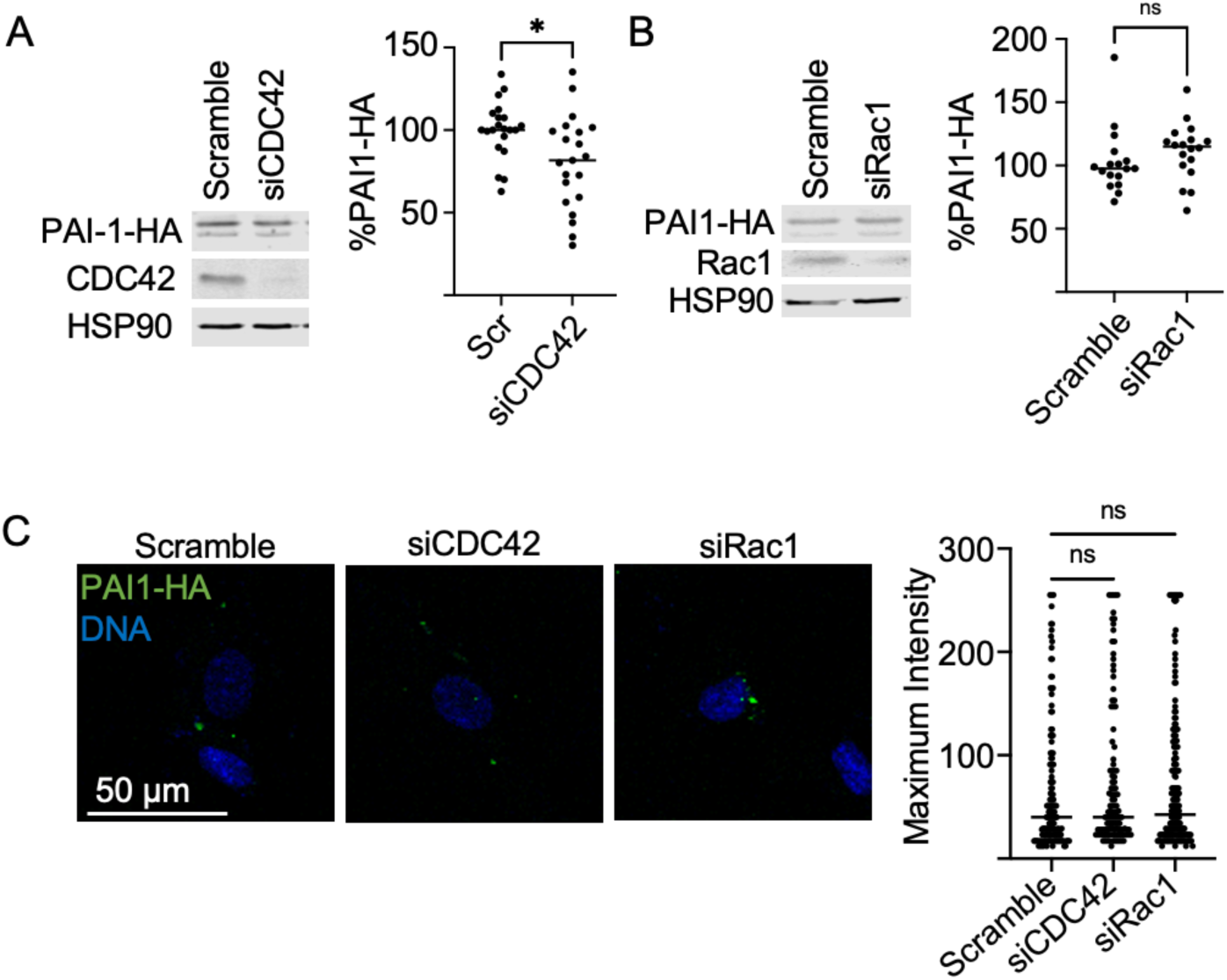
Silencing CDC42 partially reduces PAI1-HA endocytosis. Ea.hy926 cells were transfected with siRNA targeting CDC42 (A) or Rac1 (B) for 48 hours, followed by incubation with PAI1-HA spent media for two hours. Internalized PAI-1 was detected by immunoblotting toward the HA tag. Bar graph displays mean ± SEM of triplicate experiments, normalized to HSP90 loading control. C) Representative images and quantification of PAI1-HA puncta in cells treated with control siRNA or siRNA targeting CDC42 or Rac1. Graph depicts quantification of maximum intensity within each cell measured. Line depicts median. * denotes *P* <0.05. All experiments were conducted at least three separate times for reproducibility.

## Discussion

Plasminogen activator inhibitor-1 is a member of the serine protease inhibitor (SERPIN) superfamily. Originally characterized for its role in negatively regulating fibrinolysis, it is increasingly understood as a biomarker of metabolic syndrome and acute inflammatory states such as acute respiratory distress syndrome [34]. In addition to inhibiting tPA and uPA, it also inhibits other serine proteases [35]. Biological activity of intracellular PAI-1 has recently been demonstrated by several independent groups [17–19], but it is unknown how PAI-1, which has a canonical signal peptide and is secreted into the extracellular space, is endocytosed and evades lysosomal degradation. Here, we confirm that exogenous PAI1-HA added to endothelial cells in culture can be internalized and persist for several hours. Using genetic silencing and chemical inhibitors, we found that PAI1-HA internalization is stimulated by growth factors and inhibited by amiloride and that the GTPases CDC42 and Rac1 are required for internalization. These findings suggest that PAI1-HA is internalized by macropinocytosis.

Our finding that PAI1-HA is internalized via macropinocytosis is contradictory to the prior understanding that PAI-1 is endocytosed through direct interaction with LRP1 and uPAR on the cell surface. PAI-1 in its active conformation acts as a bait for its target serine protease; cleavage of its target sequence results in a conformational change that traps the serine protease [36]. Under physiologic conditions, the active conformation is unstable with a half-life of 1-2 hours [5]. Prior studies that demonstrated the role for LRP1 and uPAR have tracked the fate of the PAI-1-uPA/tPA complex [12, 37, 38]. However, the active conformation of PAI-1 that can interact with tPA or uPA is highly unstable. PAI-1 exists in its active conformation for a very short half-life and rapidly folds into its latent conformation, which is unable to bind uPA/tPA. Our recent study demonstrated that PAI1-inhibition of eNOS can occur in the latent or active conformation [18]. The conformational status was not directly studied by Bernot *et al,* but the acidic environment of the Golgi is anticipated to stabilize the active conformation of PAI1 [17]. The conformational state was not addressed in Furuya *et al* [19]. It is possible that PAI-1 bound to its target protease forms the ternary complex with uPAR and LRP1 on the cell surface, is internalized, and routed for lysosomal degradation as previously described, whereas unbound, latent PAI-1 is internalized by macropinocytosis and escapes rapid degradation. In this way, the endocytic mechanism by which PAI-1 is internalized could have differential signaling effects, allowing the endothelial cell to respond to its extracellular environment.

We are not the first to report that endothelial cells use macropinocytosis to sense and respond to extracellular signals. Endothelial cells internalize fibroblast growth factor 2 (FGF2) via a pinocytic mechanism that is dependent on Rac1 and CDC42 and inhibited by amiloride [23]. Basagiannis and colleagues found that when vascular endothelial growth factor (VEGF) receptor is internalized via macropinocytosis with VEGF, angiogenesis and migration are promoted, whereas constitutive recycling of VEGFR occurs via clathrin mediated endocytosis [25]. Moreover, microtubule activity is required for VEGF-induced endothelial cell migration [39]. Lastly, other membrane receptors such as FGFR1, transforming growth factor β, and epidermal growth factor receptor have been reported to be internalized by different mechanisms under different cellular conditions [24].

To track the fate of exogenous PAI-1, we used a construct with a C-terminal hemagglutinin tag, which could be probed with primary antibody. A limitation to this approach is that the concentration of PAI1-HA is unknown and there were batch-specific differences in concentration between experiments. Despite this variation, the percentage of cells that internalized PAI1-HA did not vary greatly from experiment to experiment, typically ∼10% after 2 hours of incubation.

It is surprising that silencing CDC42 and Rac1 did not decrease PAI1-HA uptake as visualized by our Western blotting approach. In some experiments, there was a small decrease in uptake with CDC42 silencing, but this was not statistically significant. One possible explanation is that endothelial cells have promiscuity with regards to GTPase activity and can use either GTPase depending on availability within the cell. Alternatively, silencing reduced but did not eliminate expression; small amounts of GTPase may be sufficient to support actin polymerization. Interestingly, silencing Rac1 appeared to result in a small increase in PAI1-HA by Western blotting, which could represent the increase in surface-associated PAI1-HA that could not be internalized when visualized by immunofluorescence microscopy.

PAI-1 has been assumed to contribute to disease by stabilizing fibrin networks, thereby promoting fibrosis. Increasing data suggest that PAI1’s intracellular activities may also have pathologic effects. By inhibiting furin proprotein convertase in the Golgi apparatus, PAI1 reduces insulin receptor and ADAM17 production, a proposed mechanism for a mechanistic role of PAI1 in Type 2 diabetes mellitus [17]. PAI1 appears to contribute to hypertension and fibrosis from chronic L-NAME exposure, as PAI1 knockout mice are resistant to the phenotype [40, 41], and it is reduced with a small molecule inhibitor of PAI1 [6]. Our recent finding that exogenous PAI1 inhibits endothelial nitric oxide synthase activity suggests a direct intracellular mechanism for these findings [18]. Given the multiple mechanisms by which PAI1 contributes to pathology, it is critical to better understand its biology. The studies reported here suggest an alternate mechanism by which exogenous PAI-1 can enter the intracellular space. Future studies should explore blocking this pathway blocks the intracellular activities of PAI-1.

